# Causal Lesion Evidence for Two Motor Speech Coordination Networks in the Brain

**DOI:** 10.1101/2025.06.05.658124

**Authors:** William Burns, Emma Strawderman, Steven P. Meyers, Tyler Schmidt, Kevin A. Walter, Webster H. Pilcher, Bradford Z. Mahon, Frank E. Garcea

## Abstract

Speech production is supported by sensory-to-motor transformations to coordinate activity of the larynx and orofacial muscles. Here, we show that lesions to left temporal lobe areas involved in pitch processing cause reduced neural responses when repeating sentences and when humming piano melodies in a dorsal portion of the left precentral gyrus linked to laryngeal motor control. In contrast, lesions to left inferior parietal areas involved in somatosensory processing of speech cause reduced neural responses when repeating sentences but not when humming piano melodies in a ventral portion of the left precentral gyrus linked to orofacial motor control. Analyses in neurotypical participants converge in showing that the dorsal and ventral portions of the left precentral gyrus exhibit strong functional connectivity to left temporal and inferior parietal regions, respectively. These results provide causal lesion evidence that dissociable networks underlie distinct sensory-to-motor transformations supporting laryngeal and orofacial motor control for speech production.

## Introduction

Speech production requires motor control over laryngeal and orofacial muscles to support breathing, phonation, and articulation [1–6]. Studies using functional MRI (fMRI), direct cortical stimulation, and connectome-based lesion-symptom mapping suggest that dissociable networks underlie motor control of the larynx and orofacial articulators [7–15], with separate nodes of each network in lateral portions of the left precentral gyrus. These left precentral areas are referred to as the left dorsal precentral speech area (dPCSA) and the left ventral precentral speech area (vPCSA) [16]. Adjudicating the functional role of the dPCSA and vPCSA within the broader network of brain areas supporting speech production is the focus of this fMRI investigation.

The dPCSA is situated in the dorso-lateral portion of the left precentral gyrus, anterior to the dorsal laryngeal motor cortex, and is proposed to coordinate laryngeal movements underlying prosodic elements of speech and melody [16]. For example, neural activity in the dPCSA aligns closely in time to pitch accents in speech and melodies when singing, suggesting a role in pitch coordination [17]. The dPCSA exhibits a pattern of spectro-temporal activity similar to auditory cortex–including a substantial response to vocal pitch [18], and is responsive not only during speech production but also during speech listening and pitch discrimination tasks [17, 19–22]. Moreover, there is strong resting state functional connectivity between the dPCSA and auditory regions that support pitch processing, including Heschl’s gyrus and the left superior temporal gyrus [16].

The vPCSA is situated in the ventro-lateral portion of the left precentral gyrus, anterior to the ventral laryngeal motor cortex, and is proposed to coordinate orofacial movements underlying speech articulation [16]. There is a somatotopic organization within the ventro-lateral precentral gyrus, with particular regions tuned to different orofacial and laryngeal effectors [9, 22, 23]. The vPCSA has a well-established role in generating articulatory gestures [24], and damage to this structure can be associated with apraxia of speech ([25, 26]; but see [12]). The vPCSA exhibits strong functional neural responses when correcting for experimentally induced jaw perturbations during speech production, which closely resembles responses in the anterior supramarginal gyrus, an area implicated in somatosensory processing of the jaw and lips [27, 28]. Consistent with these findings, there is strong resting state functional connectivity between the vPCSA and areas involved in somatosensory processing of speech, including the left supramarginal gyrus [16].

A dual motor speech coordination model proposes that the dPCSA interacts with auditory regions for laryngeal motor control of pitch while the vPCSA interacts with the supramarginal gyrus for orofacial motor control of articulatory gestures [16]. Here, resting state functional connectivity analyses in two independent neurotypical participant cohorts were performed to reproduce the finding that the dPCSA exhibits stronger functional connectivity to the left superior temporal gyrus than to the left anterior supramarginal gyrus, and, vice versa, the vPCSA exhibits stronger functional connectivity to the left anterior supramarginal gyrus than to the left superior temporal gyrus [16]. Next, we investigated whether these functional connectivity patterns were mirrored in patient-based analyses testing where brain lesions were associated with disruptions in functional neural responses. Sixty-six individuals with brain lesions in the preoperative phase of their neurosurgical care underwent fMRI while taking part in a sentence and melody repetition task [29]. Functional neural responses in the dPCSA and the vPCSA were extracted for sentence repetition and melody humming events (over and above sentence listening and melody listening events, respectively). Using a method referred to as Voxel-based Lesion Activity Mapping (VLAM) [30], we tested where voxelwise lesion presence was associated with weaker functional neural responses in the dPCSA and vPCSA to make causal inferences about the brain networks supporting speech production.

Given that laryngeal motor control is critical for both speech and melody humming [17], we hypothesized that participants with lesions to left Heschl’s gyrus and the left superior temporal gyrus would exhibit reduced functional neural responses in the dPCSA when repeating sentences and when humming melodies. Such findings would suggest the dPCSA interacts with regions associated with pitch processing to coordinate laryngeal movements for speech and melody production. By contrast, given that control of articulatory muscles has a more dominant role in speech relative to melody humming [17], we hypothesized that lesions to the left supramarginal gyrus would be associated with reduced functional neural responses in the vPCSA when repeating sentences but not when humming melodies. This finding would suggest that the vPCSA interacts with regions associated with somatosensory processing to coordinate articulatory movements for speech production.

## Results

### Resting State Functional Connectivity in Neurotypical Participants

Two neurotypical participant groups took part in a resting state fMRI experiment. The first group was composed of 55 relatively younger neurotypical participants, and the second group was composed of 107 relatively older neurotypical participants who were matched in age to the participants with brain lesions (see methods for details). Resting state fMRI data were used to test the hypothesis that the dPCSA would exhibit stronger functional connectivity to areas supporting pitch processing (left superior temporal gyrus), whereas the vPCSA would exhibit stronger functional connectivity to areas supporting somatosensory processing (left anterior supramarginal gyrus). The dPCSA and vPCSA regions-of-interest (ROIs) were defined using the peak coordinates reported by Hickok and colleagues (2023), which were obtained from an fMRI study of syllable perception and production conducted by Rong and colleagues (2018). For the purpose of our investigation, spherical ROIs 5 mm in diameter were drawn around the dPCSA and vPCSA peak voxels in MNI space (see Supplementary Figure 1A). Spherical ROIs 5 mm in diameter were also drawn around coordinates for the left superior temporal gyrus (STG) and left anterior supramarginal gyrus (aSMG) which were derived from Neurosynth (see Supplementary Figure 1B).

We conducted a mixed ANOVA with the within-subject factors of Seed Region (dPCSA, vPCSA) and Mask Region (STG, aSMG) and the between-subject factor of Group. Consistent with our hypotheses, there was a significant interaction between Seed Region and Mask Region (*F*(1, 159) = 134.17, p < 0.001, ƞ² = 0.48). Post-hoc *t*-test revealed that the dPCSA exhibited stronger functional connectivity to the left STG than to the left aSMG (younger group: bar 1 > bar 2, *t*(54) = 3.12, *p* < 0.01; age-matched group: bar 5 > bar 6, *t*(105) = 3.30, *p* < 0.001), and, vice versa, the vPCSA exhibited stronger functional connectivity to the left aSMG than to the left STG (younger group: bar 4 > bar 3, *t*(54) = 7.00, *p* < 0.001; age-matched group: bar 8 > bar 7, *t*(105) = 6.83, *p* < 0.001; see Figure 1).

**Figure 1.**
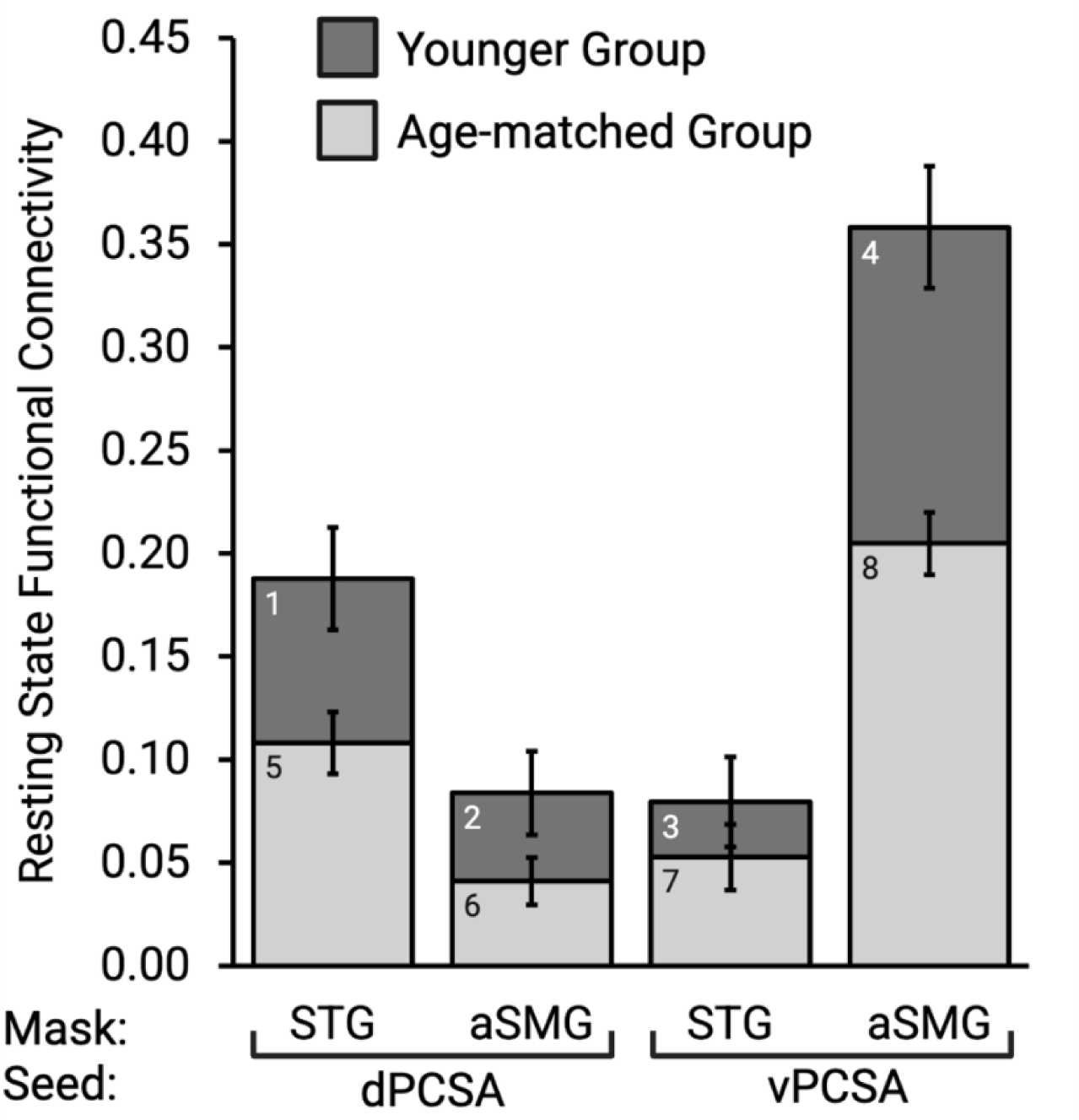
Resting state functional connectivity analysis of the dPCSA and vPCSA. Across both neurotypical participant groups, the dPCSA exhibited stronger functional connectivity to the left superior temporal gyrus (STG), whereas the vPCSA exhibited stronger functional connectivity to the left anterior supramarginal gyrus (aSMG). Error bars represent standard error of the mean computed across participants.

### Analysis of Functional MRI Responses in Neurosurgical Participants

Next, we studied a series of 66 consecutively enrolled patient participants in the preoperative phase of their neurosurgical care at the University of Rochester. Participants completed a melody and sentence repetition fMRI task, in which they listened to 3-second clips of sentences and piano melodies and reproduced the stimuli after a delay period of several seconds. This design feature allowed us to model the sentence and melody listening phases of the conditions separately from the sentence and melody reproduction phases (for precedent, see [29]). Some participants completed a version of the task which included repetition and completion of short arithmetic questions, which was not analyzed here, as it is not germane to the focus of the present investigation. The areas of greatest lesion overlap among the 66 participants were in the left hemisphere, and included the temporal pole, hippocampus, insula, pre- and post-central gyri, the superior frontal gyrus, and temporo-parietal regions including the superior temporal gyrus and the supramarginal gyrus (see Supplementary Table 1 for lesion etiology and demographic variables for all participants, and Supplementary Figure 2 for a lesion overlap map).

Within the dPCSA and vPCSA ROIs, we extracted contrast-weighted *t*-values separately for sentence and melody listening phases as well as for sentence and melody reproduction phases. We then conducted a repeated-measures ANOVA with the factors Region (dPCSA, vPCSA), Stimulus Condition (Sentences, Melodies) and Task Phase (Listen, Reproduce) to test the hypothesis that the dPCSA and vPCSA ROIs would exhibit robust functional neural responses in the listening and reproduction phases of both conditions. Consistent with that hypothesis, there was no main effect of Region, Stimulus Condition, or Task Phase, nor any significant two-way or three-way interaction effects. As is seen in Figure 2, there were robust responses for the listening and reproduction phases of both conditions in both the dPCSA and vPCSA.

**Figure 2.**
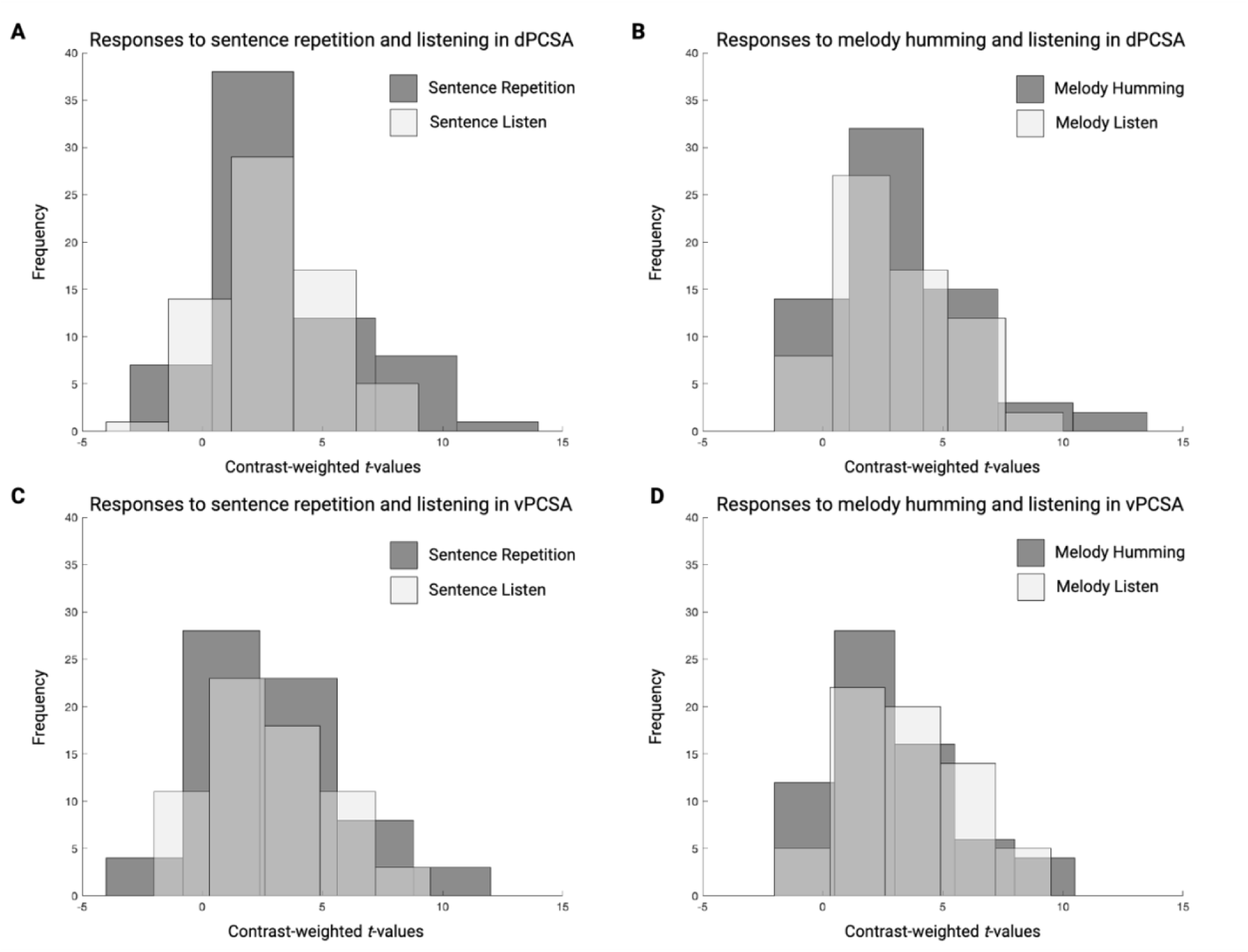
ROI analysis of condition and phase in the dorsal and ventral precentral speech areas. Light bars represent the distribution of contrast-weighted *t*-values during the listening phase, whereas dark bars represent the distribution of contrast-weighted *t*-values during the production phase. Across both dPCSA (A, B) and vPCSA (C, D) ROIs, there was considerable overlap in the distribution of values for the listening and reproduction phases of each condition, which explains the lack of significant main effects or interaction effects in the repeated-measures ANOVA.

### Voxel-based Lesion-Activity Mapping (VLAM) of Sentence Repetition in the dPCSA and vPCSA

VLAM is a technique in which functional neural responses in a ROI are used to predict variance in voxel-wise lesion presence throughout the brain. Using VLAM, we inspected the lesion sites associated with reduced functional neural responses in the dPCSA and vPCSA when producing sentences, and separately, when humming melodies. To that end, we extracted contrast-weighted *t*-values from the dPCSA and vPCSA ROIs using the contrasts ‘Sentence Repetition > Sentence Listening’ and ‘Melody Humming > Melody Listening’ in every participant. These values served as the independent variables in the VLAM analyses.

In the first VLAM analysis, we entered the contrast-weighted *t*-values for ‘Sentence Repetition > Sentence Listening’ in the dPCSA as the independent variable, and tested where lesion presence (dependent variable) was inversely related to the amplitude of functional neural responses. Our first hypothesis was that participants with lesions to left Heschl’s gyrus and the left superior temporal gyrus would exhibit reduced functional neural responses in the dPCSA when repeating sentences. Consistent with this hypothesis, functional neural responses in the dPCSA were reduced in association with lesion presence in left Heschl’s gyrus, the left planum temporale, and the left superior temporal gyrus. This cluster included the posterior portions of the superior temporal gyrus, the ventral supramarginal gyrus and angular gyrus, the dorsal posterior middle temporal gyrus, and the pre- and post-central gyri (see Figure 3, red-to-yellow, and Table 1A).

**Figure 3.**
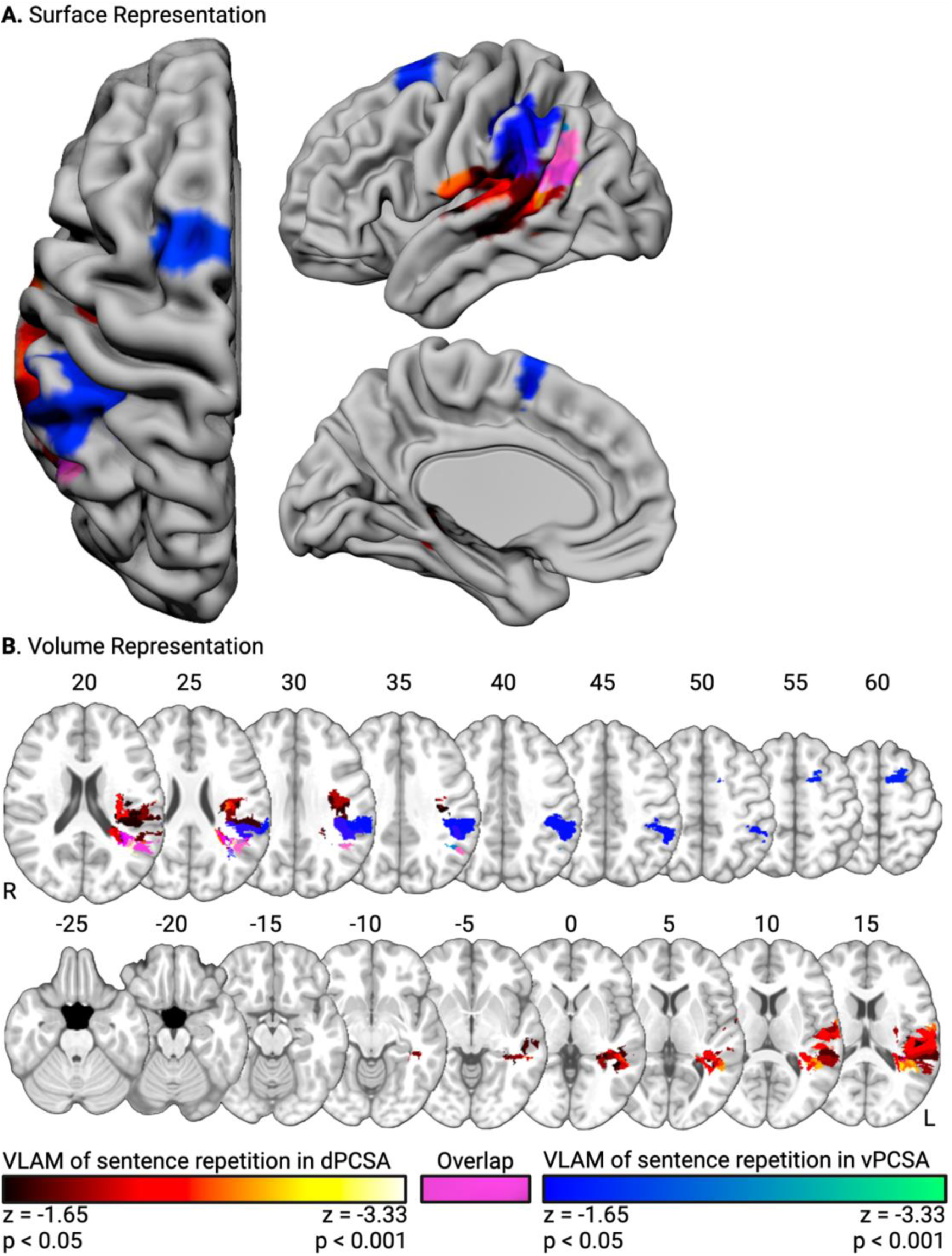
Lesion sites inversely associated with functional neural responses for sentence repetition. Surface-based (A) and volumetric (B; MNI Z coordinate listed above axial slices) renderings of lesion sites associated with reduced functional neural responses in the dPCSA (red- to-yellow) and the vPCSA (blue-to-green). Common lesion sites are rendered in magenta.

**Table 1.** Regions and MNI coordinates of voxels identified in the VLAM analysis of sentence repetition in (A) the dPCSA and (B) the vPCSA. Regions of overlap across analyses (C).

| Region Name | Region Number | Peak MNI Coordinate |  |  | Peak Z-Score | Percent of Region with Significant Voxels |
| --- | --- | --- | --- | --- | --- | --- |
|  |  | X | Y | Z |  |  |
| A. VLAM of sentence repetition in the dPCSA |  |  |  |  |  |  |
| Angular gyrus | 41 | -39 | -57 | 19 | -3.35 | 17.80 |
| Temporo-parieto-occipital portion of the left middle temporal gyrus | 25 | -51 | -47 | 8 | -2.75 | 13.88 |
| Posterior supramarginal gyrus | 39 | -49 | -48 | 12 | -2.75 | 4.41 |
| Planum temporale | 91 | -42 | -40 | 13 | -2.74 | 51.95 |
| Precentral gyrus | 13 | -63 | -3 | 11 | -2.56 | 25.96 |
| Postcentral gyrus | 33 | -63 | -11 | 16 | -2.56 | 0.80 |
| Central operculum | 83 | -59 | -14 | 17 | -2.56 | 0.03 |
| Posterior middle temporal gyrus | 23 | -50 | -39 | -1 | -2.48 | 4.93 |
| Posterior superior temporal gyrus | 19 | -68 | -23 | 12 | -2.42 | 74.60 |
| Parietal operculum | 85 | -49 | -30 | 16 | -2.42 | 26.44 |
| Heschl's gyrus | 89 | -44 | -26 | 13 | -2.42 | 13.77 |
| Insular cortex | 3 | -35 | -21 | 18 | -2.25 | 0.64 |
| Posterior fusiform gyrus | 75 | -36 | -40 | -11 | -1.98 | 0.04 |
| Anterior supramarginal gyrus | 37 | -65 | -26 | 19 | -1.92 | 0.79 |
| B. VLAM of sentence repetition the vPCSA |  |  |  |  |  |  |
| Angular gyrus | 41 | -39 | -59 | 23 | -2.69 | 15.81 |
| Posterior supramarginal gyrus | 39 | -51 | -51 | 26 | -2.40 | 14.85 |
| Superior frontal gyrus | 5 | -22 | 12 | 62 | -1.95 | 5.83 |
| Supplementary motor area | 51 | -8 | 2 | 60 | -1.90 | 2.64 |
| Anterior supramarginal gyrus | 37 | -59 | -37 | 29 | -1.76 | 40.36 |
| Parietal operculum | 85 | -51 | -37 | 26 | -1.76 | 17.45 |
| Postcentral gyrus | 33 | -41 | -32 | 44 | -1.73 | 1.33 |
| Superior parietal lobe | 35 | -43 | -40 | 50 | -1.73 | 0.77 |
| C. Overlap across analyses |  |  |  |  |  |  |
| Angular gyrus | 41 | -48 | -56 | 25 | - | 15.36 |
| Parietal operculum | 85 | -51 | -37 | 26 | - | 14.26 |
| Posterior supramarginal gyrus | 39 | -54 | -44 | 29 | - | 2.99 |
| Anterior supramarginal gyrus | 37 | -59 | -37 | 29 | - | 0.38 |

Our second hypothesis was that participants with lesions to the left anterior supramarginal gyrus would exhibit reduced functional neural responses in the vPCSA when repeating sentences. Consistent with this hypothesis, functional neural responses in the vPCSA were reduced in association with lesion presence in the left anterior supramarginal gyrus. This cluster included the angular gyrus, the parietal operculum, and postcentral gyrus. A second cluster was identified in the left superior frontal gyrus and left pre-supplementary motor area (see Figure 3, blue-to-green, and Table 1B). Across the analyses, overlapping voxels were identified in the angular gyrus and white matter undercutting the left supramarginal gyrus and the left parietal operculum (Figure 3, magenta, and Table 1C).

### VLAM of Melody Humming in the Dorsal and Ventral Precentral Speech Areas

Contrast-weighted *t*-values for ‘Melody Humming > Melody Listening’ in the dPCSA and vPCSA served as independent variables in two subsequent VLAM analyses. The third hypothesis was that participants with lesions to left Heschl’s gyrus and the left superior temporal gyrus would exhibit reduced functional neural responses in the dPCSA when humming melodies. Consistent with this hypothesis, functional neural responses in the dPCSA were reduced in association with lesions to left Heschl’s gyrus, the left planum temporale, and the left superior temporal gyrus. This cluster extended into posterior portions of the left superior and middle temporal gyri, and the white matter undercutting the supramarginal gyrus and angular gyrus. An adjacent lesion cluster included in a ventral portion of the left pre- and post-central gyri and parietal operculum (see Figure 4, red-to-yellow, and Table 2A). The final VLAM analysis tested whether lesions to areas supporting somatosensory processing in the left anterior supramarginal gyrus were associated with weaker functional neural responses in the vPCSA when humming melodies. We hypothesized that they would not, and consistent with that prediction, we found that lesions to the left anterior temporal lobe, the left middle and superior frontal gyri, and a small portion of the left pre- and post-central gyri–but not the left anterior supramarginal gyrus–were associated with reduced functional neural responses in the vPCSA when participants hummed melodies (see Figure 4, blue-to-green, and Table 2B). No overlapping voxels were identified between the two VLAM analyses of melody humming.

**Figure 4.**
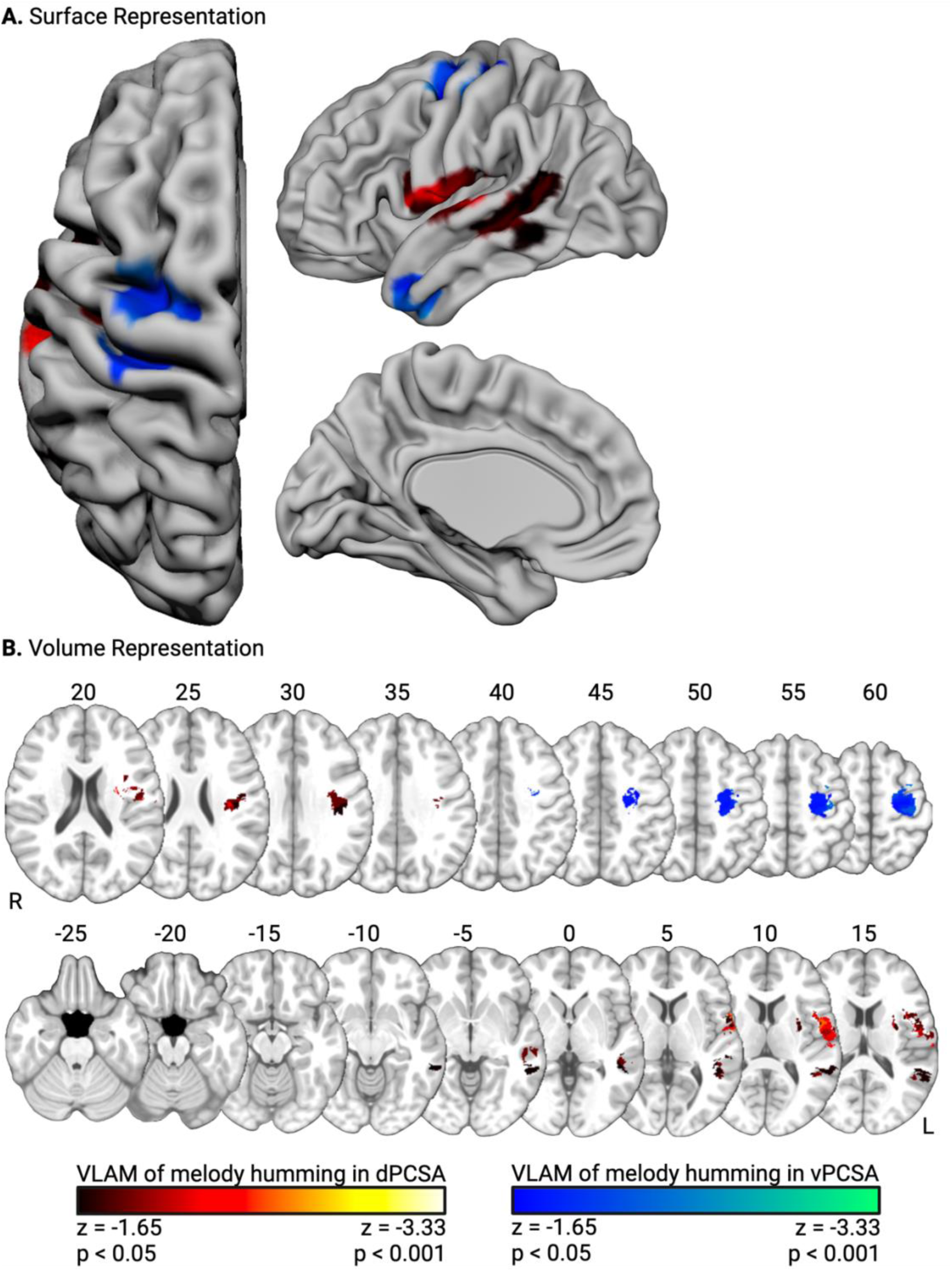
Lesion sites inversely associated with functional neural responses for melody humming. Surface-based (A) and volumetric (B; MNI Z coordinate listed above axial slices) renderings of lesion sites associated with reduced functional neural responses in the dPCSA (red-to-yellow) and the vPCSA (blue-to-green).

**Table 2.** Regions and MNI coordinates of voxels identified in the VLAM analysis of melody humming in (A) the dPCSA and (B) the vPCSA.

| Region Name | Region Number | Peak MNI Coordinate |  |  | Peak Z-Score | Percent of Region with Significant Voxels |
| --- | --- | --- | --- | --- | --- | --- |
|  |  | X | Y | Z |  |  |
| A. VLAM of melody humming in the dPCSA |  |  |  |  |  |  |
| Precentral gyrus | 13 | -52 | 3 | 13 | -2.91 | 1.71 |
| Inferior frontal gyrus | 11 | -59 | 10 | 5 | -2.57 | 1.05 |
| Central operculum | 83 | -53 | -4 | 10 | -2.57 | 21.46 |
| Heschl's gyrus | 89 | -53 | -13 | 8 | -2.57 | 2.16 |
| Planum temporale | 91 | -60 | -14 | 8 | -2.57 | 2.41 |
| Postcentral gyrus | 33 | -66 | -14 | 14 | -2.25 | 0.61 |
| Posterior superior temporal gyrus | 19 | -63 | -33 | 8 | -2.01 | 5.96 |
| Posterior supramarginal gyrus | 39 | -48 | -51 | 16 | -1.97 | 5.90 |
| Angular gyrus | 41 | -49 | -52 | 16 | -1.97 | 2.29 |
| Temporo-parieto-occipital portion of the left middle temporal gyrus | 25 | -51 | -47 | 8 | -1.90 | 5.00 |
| Posterior middle temporal gyrus | 23 | -54 | -31 | -3 | -1.87 | 6.38 |
| Anterior superior temporal gyrus | 17 | -62 | -2 | 2 | -1.80 | 0.94 |
| Planum polare | 87 | -58 | -5 | 4 | -1.80 | 0.99 |
| B. VLAM of melody humming in the vPCSA |  |  |  |  |  |  |
| Temporal pole | 15 | -50 | 8 | -31 | -2.86 | 5.02 |
| Middle frontal gyrus | 7 | -33 | 0 | 59 | -2.28 | 0.54 |
| Precentral gyrus | 13 | -35 | -2 | 55 | -2.28 | 7.95 |
| Anterior middle temporal gyrus | 21 | -48 | 0 | -31 | -2.20 | 0.03 |
| Anterior inferior temporal gyrus | 27 | -45 | 1 | -36 | -2.20 | 0.23 |
| Postcentral gyrus | 33 | -35 | -25 | 60 | -2.18 | 0.84 |
| Superior frontal gyrus | 5 | -24 | -6 | 63 | -2.06 | 1.74 |

### Comparisons Between VLAM Analyses of Sentence Repetition and Melody Humming

A core prediction of the dual motor speech coordination model is that the dPCSA interacts with auditory regions for laryngeal motor control of pitch. Consistent with that prediction, we identified overlapping lesion sites in the left hemisphere associated with reduced functional neural responses for speech repetition *and* melody humming in the dPCSA. Those sites included Heschl’s gyrus, the planum temporale, and the superior temporal gyrus; a second site included the white matter undercutting the supramarginal gyrus, the angular gyrus, and the posterior middle temporal gyrus; and a third cluster included a ventral portion of the central operculum, and pre- and post-central gyri (see Figure 5 and Table 3). A second core prediction is that the vPCSA interacts with the supramarginal gyrus for orofacial motor control of articulatory gestures. Consistent with that prediction, we found that lesions to the left supramarginal gyrus were associated with reduced functional neural responses in the vPCSA for sentence repetition but not for melody humming (see Figures 3 & 4). In subsequent analyses, we confirmed that the magnitude of the association between anterior supramarginal gyrus lesion damage and functional neural responses in the vPCSA for sentence repetition was statistically stronger than the magnitude of the association between anterior supramarginal gyrus lesion damage and functional neural responses in the vPCSA for melody repetition (see Supplemental Figure 3 and Supplementary Table 2).

**Figure 5.**
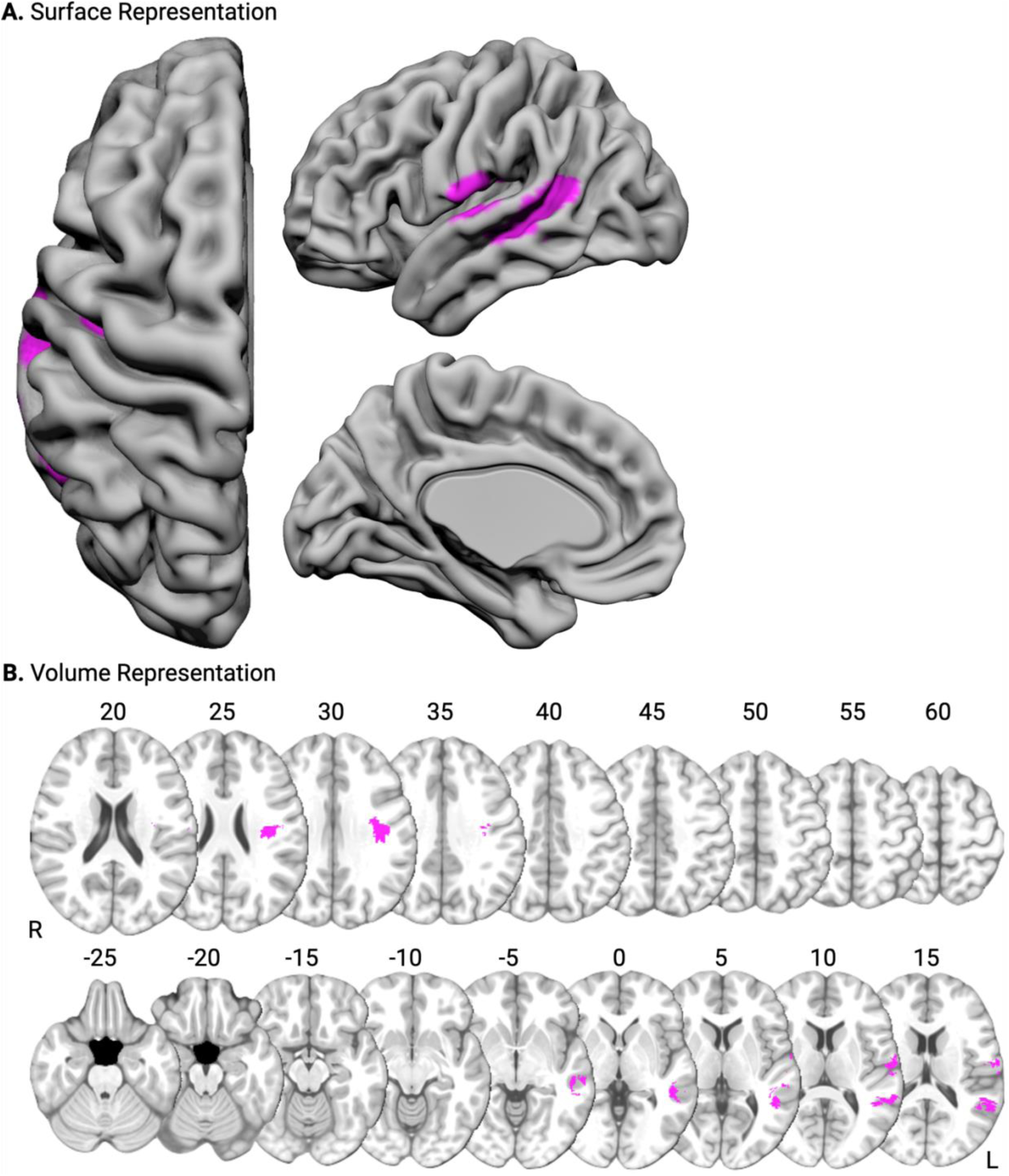
Overlap between VLAM of sentence repetition and melody humming in the dPCSA. Surface-based (A) and volumetric (B; MNI Z coordinate listed above axial slices) renderings of the overlapping lesion sites identified in the VLAM analysis of sentence repetition and melody humming in the dPCSA.

**Table 3.** Overlapping regions identified in the VLAM analysis of sentence repetition and melody humming in the dPCSA.

| Region Name | Region Number | Peak MNI Coordinate |  |  | Percent of Region with Significant Voxels |
| --- | --- | --- | --- | --- | --- |
|  |  | X | Y | Z |  |
| Central operculum | 83 | -59 | -13 | 13 | 7.47 |
| Posterior superior temporal gyrus | 19 | -60 | -35 | 6 | 5.96 |
| Posterior supramarginal gyrus | 39 | -57 | -46 | 13 | 5.90 |
| Posterior middle temporal gyrus | 23 | -54 | -32 | -3 | 4.15 |
| Temporo-parieto-occipital portion of the left middle temporal gyrus | 25 | -50 | -47 | 5 | 3.36 |
| Angular gyrus | 41 | -57 | -52 | 16 | 2.29 |
| Planum temporale | 91 | -60 | -19 | 9 | 2.18 |
| Heschl's gyrus | 89 | -55 | -18 | 9 | 1.24 |
| Postcentral gyrus | 33 | -62 | -12 | 17 | 0.61 |
| Precentral gyrus | 13 | -63 | -3 | 11 | 0.03 |

### General Discussion

Motor control over laryngeal and orofacial muscles is critical to enable breathing, phonation, and articulation for speech production. The objective of this report was to identify the underlying functional brain networks supporting laryngeal and orofacial motor control from the perspective of the dorsal (dPCSA) and ventral precentral speech areas (vPCSA). We first demonstrated that the dPCSA exhibited differentially stronger resting state functional connectivity to the left superior temporal gyrus, whereas the vPCSA exhibited differentially stronger resting state functional connectivity to the left anterior supramarginal gyrus. This finding replicates prior work [16] and was identified in two independent neurotypical cohorts. Next, we showed that functional neural responses in the dPCSA and vPCSA were attenuated in association with lesion damage to dissociable temporal and parietal areas: Lesions to left Heschl’s gyrus and the adjacent left superior temporal gyrus led to reduced functional neural responses in the dPCSA when repeating sentences *and* humming melodies, whereas lesions to the left anterior supramarginal gyrus led to reduced functional neural responses in the vPCSA *only* when repeating sentences. Such findings indicate that lesions can disrupt processing distal to the site of injury in a task-specific manner, a phenomenon Price and colleagues (2001) have referred to as dynamic diaschisis (for discussion, see [32, 33]). These results provide causal lesion evidence in support of a model whereby the dPCSA and vPCSA are part of separable networks interacting with early auditory and somatosensory regions, respectively, to coordinate motor activity of the larynx and orofacial muscles for speech production, respectively.

State feedback control models posit that sensory and motor control systems interact to coordinate muscle activity of the vocal tract and orofacial articulators during speech production [34, 35]. The measured sensory consequences of speech movements in the form of auditory or somatosensory feedback are compared to feedforward predictions of the auditory and somatosensory consequences or targets of the planned motor action. In the context of our task, pitch representations of the sentence and melody stimuli are the auditory targets, and somatosensory representations of the articulatory gestures are the somatosensory targets. When reproducing a sentence, differences between the predicted sensory targets and real-time sensory feedback of the speech movements generate an error signal, which is used to update the motor commands to aid in the generation of the intended speech. This mechanism allows for error-correction, motor learning, and sensory adaptation (e.g., see [36–38]. For example, experimentally induced shifts in perceived auditory feedback result in compensatory changes in produced pitch [37, 38], and introducing auditory feedback delays result in disrupted speech fluency [39]. In the somatosensory domain, unexpected robotic perturbations to the jaw during speech production result in gradual adaptation of jaw movements [36].

Attenuations of functional neural responses in the dPCSA and vPCSA could represent (1) disruptions in the generation of feedforward predictions, or (2) degraded processing of feedback to generate error signals. Although future work is needed to adjudicate between these possible mechanisms, converging evidence from our functional connectivity and VLAM analyses suggest the dPCSA interacts with regions associated with auditory processing to coordinate laryngeal movements during the generation of both speech and melody humming, whereas the vPCSA interacts with regions associated with somatosensory processing to coordinate articulatory movements during the generation of speech. These functional networks are consistent with regions of the brain implicated in sensorimotor feedback control. For example, auditory perturbation paradigms probing auditory-to-motor transformations implicate both left dorsal precentral gyrus and superior temporal gyrus [21], while speech perturbation paradigms probing somatosensory-to-motor transformations implicate the ventral precentral gyrus and supramarginal gyrus [27, 40].

It’s important to recognize that somatosensory feedback is not irrelevant for the reproduction of melodies, as the relative strength of somatosensory feedback may vary depending on task demands [41]. For instance, if participants were instructed to sing melodies which emphasize orofacial articulation (e.g., “do-re-mi-re-do”; c.f., [17]) rather than hum melodies, lesions to the anterior supramarginal gyrus may alter somatosensory feedback, resulting in reduced functional neural responses in the vPCSA. Differences in somatosensory feedback weight may thus depend on the articulatory gestures cued by the task, with a greater emphasis placed on somatosensory feedback of vowels that involve lingual contact [42]. Future studies manipulating the recruitment of orofacial articulation during speech and melody production are needed to better understand the salience of somatosensory feedback—and underlying neuroanatomic substrates—given processing demands imposed by the task.

There was partial lesion overlap between VLAM analyses of sentence repetition in the left angular gyrus, left parietal operculum, and left posterior supramarginal gyrus–suggesting these areas may support multimodal sensory integration. One hypothesis is that somatosensory processing may provide input to auditory systems about timing onset and rate of articulation to refine auditory-to-motor transformations, and, vice versa, the auditory system may refine somatosensory processing to inform the somatosensory-to-motor mapping processes underlying orofacial gesture production [34]. A second hypothesis is that these areas arbitrate the relative contributions of auditory and somatosensory error signals in a given task to optimize task-specific feedback correction [43]. For example, in one speech repetition study where somatosensory and auditory feedback were simultaneously altered, there was typically a preferential reliance on one feedback system–with considerable variation across participants–implying the existence of flexible feedback control systems to arbitrate sensory error signals [44].

Lesions to the ventral and lateral precentral gyrus, near the vPCSA, were associated with reduced functional neural responses to sentence repetition and melody humming in the dPCSA. We suspect this finding is related to tight coupling of laryngeal motor control of the vPCSA and dPCSA [45]. The ventral and lateral precentral gyrus has been hypothesized to coordinate the intrinsic muscles of the larynx for rapid tension (i.e. adduction) and relaxation (i.e. abduction) of the vocal folds at the onset and offset of a vocalization [17, 46]. The dorsal and lateral precentral gyrus, on the other hand, may coordinate the extrinsic muscles of the larynx and the cricothyroid muscle to stretch the vocal cords for fine-tuned pitch modulation [17, 46]. Thus, without appropriate intrinsic laryngeal muscle control following damage to the ventral and lateral precentral gyrus, the dorsal and lateral precentral gyrus may be less effective in coordinating extrinsic laryngeal muscles for fine-tuned pitch modulation.

One lesion site found only in the VLAM analysis of sentence repetition in the vPCSA was the pre-supplementary motor area (pre-SMA)^1^. Whereas the SMA-proper is involved in motor initiation and speech articulation [47], the pre-SMA, located anterior to the SMA, is implicated in sequencing of phonological representations for articulation post lexical selection [48]. Our proposal is that when coordinating activity of the orofacial articulators for production, modulatory feedback from the pre-SMA via the frontal aslant tract [49–52] enables the planning and sequencing of phonological representations for sentence repetition. Lesions to the pre-SMA disrupt these functional interactions, resulting in a computation-specific diaschisis to the somatosensory-to-motor coordination system but not the auditory-to-motor coordination system. It will be important for future research to investigate whether the sensory-to-motor coordination functions enabled by the dorsal and ventral speech networks are bilaterally represented. Whereas meta-analyses of speech arrest errors identify dorsal and ventral foci in the left precentral gyrus [14], direct electrical stimulation to the right precentral gyrus can also lead to speech errors. For example, Belkhir and colleagues (2021) found that direct electrical stimulation to the right dorsal precentral gyrus during awake brain mapping led to guttural vocalizations and dysfluent speech that could not be resolved until after the discontinuation of the direct electrical stimulation. Interestingly, when adjacent areas were stimulated, the patient initiated vocal responses without hesitation and did not produce the guttural vocalizations. One possibility is that motor control of the larynx is bilaterally represented, whereas the integration functions subserving auditory-to-motor mapping for feedback control are left-lateralized. Whether the same functional principles apply to the somatosensory-to-motor mapping processes underlying articulatory gesture production remains an open question for future studies.

In conclusion, we provide causal lesion evidence in support of a dual motor speech coordination model, highlighting the dissociable networks underlying auditory-to-motor and somatosensory-to-motor transformations in support of sentence repetition and melody humming. Functional connectivity results in neurotypical participants complement the VLAM analyses by demonstrating that the distal areas exhibiting strong functional connectivity to the dPCSA (left superior temporal gyrus) and vPCSA (left anterior supramarginal gyrus) were the same areas that, when lesioned, led to reductions in functional neural responses in the dPCSA and vPCSA. These results suggest that damage to auditory and somatosensory systems results in computation-specific attenuation upon relatively “upstream” coordination processes mediating motor control of the larynx and orofacial articulators for speech production.

## Methods

### Participants

Seventy-four individuals in the preoperative phase of their neurosurgical care participated in this study. Participants were between the ages of 18-80 and had normal or corrected-to-normal vision (36 females; mean age 45.4 y; SD 17.0). Eight participants’ data were discarded for the following reasons: Five had no identifiable lesion, one had a prior neurosurgery, one had excessive motion artifact, and one did not complete at least 2 runs of the experiment. Of the remaining 66 participants, 48 had left hemisphere lesions and 18 had right hemisphere lesions (see Supplemental Figure 2 for lesion overlap, and Supplemental Table 1 for demographic variables). These participants took part in the study in exchange for payment and gave written informed consent in accordance with the University of Rochester Research Subjects Review Board.

A group of 55 neurotypical adults (29 females; mean age 22.3 y; SD 6.0) who were younger than the neurosurgical cohort (t(127) = 9.45, p < 0.001) participated in resting state fMRI as part of a project investigating object representations in the ventral visual pathway [54–61]. This study took place using the same MRI scanner as the neurosurgical cohort. All participants took part in the study in exchange for payment and gave written informed consent in accordance with the University of Rochester Research Subjects Review Board.

An additional 107 neurotypical adults (63 females; mean age 49.2 y; SD 15.9) who were not different in age from the neurosurgical cohort (t(179) = 1.53, p = 0.13) took part in resting state fMRI as part of a clinical evaluation for subjective hearing loss, tinnitus, and/or headache. The MRI examinations were evaluated by a neuro-radiologist (co-author SPM), who verified there were no signs of structural abnormality in these patients. All protected health information was removed from the clinical MRI images prior to data analysis. This retrospective research study was approved by the University of Rochester Research Subjects Review Board.

### Sentence and Melody Repetition fMRI Task

Participants in the neurosurgical cohort took part in two scanning sessions. The first session consisted of a T1 anatomical scan, an object processing category localizer experiment (data not analyzed herein), resting state fMRI (data not analyzed herein), and a diffusion tensor imaging scan (data not analyzed herein). The 55 younger neurotypical participants also took part in this fMRI session. The second fMRI session consisted of 2-to-8 runs of a sentence and melody repetition experiment [29], along with a motor localizer experiment (data not analyzed herein). fMRI data collection occurred over an 8-year period. Stimulus presentation was controlled with ‘A Simple Framework’ [62] written in MATLAB using the Psychophysics Toolbox [63], with PsychoPy [64], or with custom stimulus presentation software. Participants viewed stimuli binocularly through a mirror attached to the head coil adjusted to allow foveal viewing of a back-projected monitor (spatial resolution = 1400 × 1050 pixels; temporal resolution = 120 Hz). Several updates were introduced over the years, resulting in two versions of the task.

In version A of the task, participants listened to auditory stimuli and were cued to reproduce the stimulus by repeating it aloud. The stimuli consisted of 3-second clips of piano melodies made available from the Montreal Battery for the Evaluation of Amusia [65], as well as 3-second spoken sentences adapted from subtest 12 of the Psycholinguistic Assessment of Language Processing in Aphasia [66]. Each trial began with a centrally presented oval, followed by a stimulus that was presented auditorily for 3 seconds; a rehearsal period (jittered to be 12 to 20 seconds in length) followed the go cue. During the rehearsal period, participants were instructed to maintain the auditory stimulus in memory until the word ‘GO’ was presented; upon the presentation of the ‘GO’ stimulus, participants were instructed to hum or repeat the previously presented melody or sentence, respectively. Trials were interspersed with 16-second fixation periods in which a centrally presented oval was presented. Five trials of piano melodies and five trials of sentences were presented within a run, with the constraint that the two conditions were counterbalanced across runs (e.g., Subject 1, Run 1: Melody-Sentence-Melody-Sentence; Subject 1, Run 2: Sentence-Melody-Sentence-Melody).

In version B, the same piano melody stimuli were included. The sentence stimuli consisted of 16 three-second sentences pertaining to people and objects. Sentences were constructed such that the final word was not included in the recording (e.g., “the boy stopped to tie his …”). All sentences maintained high cloze probability (> 0.92) using norms provided by Peelle and colleagues [67], ensuring the final word of each sentence was constrained by the sentence context. When cued to reproduce the stimulus, participants were instructed to repeat the sentence and fill in the blank (e.g., “the boy stopped to tie his shoes”). A math condition was also included that maintained an identical task demand as the sentence stimuli. Sixteen three-second recordings of addition, subtraction, multiplication, and division equations were created for participants to listen to, rehearse, and solve (e.g., 14 + 2 = 16). Before the experiment proper, all participants were familiarized with the stimuli and were asked to hum the melody, to repeat and fill in the final word of each sentence, and to solve each math equation. This preparation phase ensured the participant could complete the task in a timely manner. We did not model the math condition in the data analysis herein, as that condition is not germane to the aims of this project.

Each trial began with a centrally presented fixation cross for 8 seconds, followed by a stimulus that was presented auditorily for 3 seconds; a jittered rehearsal period (3 to 9 seconds) followed the go cue. During the rehearsal period, participants were instructed to maintain the previously presented stimulus in memory until a beep sound was presented; upon the presentation of the beep stimulus, participants were to reproduce the previously presented melody, sentence, or math equation. Trials were interspersed with 8-second fixation periods in which a centrally presented fixation cross was presented. Four trials of piano melodies, 4 trials of sentences, and 4 trials of math equations were presented within a run, with the constraint that each condition did not repeat over 2 successive trials. After completing 12 trials (1 run), each condition was presented 4 times.

### MRI Parameters

Whole-brain BOLD imaging was conducted on a 3-Tesla Siemens MAGNETOM Trio scanner with a 20, 32, or 64-channel head coil located at the Center for Advanced Brain Imaging and Neurophysiology. High-resolution structural T1 contrast images were acquired using a magnetization-prepared rapid gradient echo (MP-RAGE) pulse sequence at the start of each participant’s scanning session (TR = 2530 ms, TE = 3.44 ms, flip angle = 7°, FOV = 256 mm, matrix = 256 × 256, 192 1 x 1 x 1 mm^3^ sagittal left-to-right slices). An echo-planar imaging pulse sequence was used for T2* contrast. In the cohort of 55 non-age-matched neurotypical participants, the resting state acquisition parameters were: TR = 2000ms, TE = 30ms, flip angle = 90°, FOV = 256 × 256 mm, matrix = 64 × 64, 30 inferior-to-superior axial slices, voxel size = 4 x 4 x 4 mm. In 32 of the neurosurgical participants, the sentence and melody repetition task acquisition parameters were: TR = 2200 ms, TE = 30 ms, flip angle = 90°, FOV = 256 x 256 mm, matrix = 64 x 64, 33 inferior-to-superior axial slices, voxel size = 4 x 4 x 4 mm^3^. In the remaining 34 neurosurgical participants, the acquisition parameters were: TR = 2200 ms, TE = 30 ms, flip angle = 70°, FOV = 256 x 256 mm, matrix = 128 x 128, 90 inferior-to-superior axial slices, voxel size = 2 x 2 x 2 mm^3^. Across all participants, the first 4 volumes of each run were discarded to allow for signal equilibration (4 volumes dropped prior to image acquisition).

The 107 age-matched neurotypical participants were scanned on a 3-Tesla Siemens SKYRA scanner with a 20-channel head coil. Each scanning session consisted of a T1 anatomical scan and 1 run of resting state fMRI. High-resolution structural T1 contrast sagittal images were acquired using an MP-RAGE pulse sequence at the start of the session (TR = 1200 or 1310 ms; TE = 2.29 or 2.33 ms; flip angle = 8 or 9°; matrix = 256 × 256, 1 x 1 x 1 mm^3^). Whole brain BOLD imaging was acquired using a Gradient Echo Planar Imaging (GE-EPI) sequence (TR = 2170– 2890 ms; TE = 30 ms; flip angle = 90°; resolution = 2 x 2 x 2 mm^3^).

### Lesion Identification

Lesions were identified by segmenting healthy tissue from the damaged tissue visible on the native T1 image. Lesioned voxels consisting of both grey and white matter were assigned a value of 1, and preserved voxels were assigned a value of 0 using ITK-SNAP software [68]. All participants provided written consent for the research team to access T1 and T2 structural imaging acquired as part of their clinical care. Lesion drawings were then reviewed by neuro-radiologist S.P. Meyers, who was at the time naïve to the aims of the project, and who had access to T1, T2, and FLAIR imaging (with and without contrast). Lesion renderings were visually inspected by the neuro-radiologist and corrected by study team members when the drawings did not accurately capture the boundaries of the lesion. Lesion renderings in the native T1 space were entered as cost function masks [69] during anatomical and functional data pre-processing. This feature ensures that the anatomical data could be normalized to Montreal Neurological Institute (MNI) space while minimizing error introduced by a lesion. The corrected lesion drawings were then normalized to MNI space using the transformation matrix that normalized the native T1 image into MNI space (using the ANTs toolbox). Finally, the accuracy of the lesions in MNI space were reviewed and confirmed, using anatomical landmarks as reference to ensure no distortions were introduced during normalization (e.g., see [70–73]).

### Anatomical Data Preprocessing

Each participant’s T1-weighted (T1w) images was corrected for intensity non-uniformity with ‘N4BiasFieldCorrection’ (ANTs), and used as T1w-reference throughout the workflow. The T1w-reference was then skull-stripped with the ‘antsBrainExtraction.sh’ workflow (from ANTs, within NiPype), using OASIS30ANTs as target template. Brain tissue segmentation of cerebrospinal fluid, white-matter, and gray-matter was performed on the brain-extracted T1w using ‘fast’ (FSL). Volume-based spatial normalization to standard space (MNI152nLin2009cAsym) was performed through nonlinear registration with ‘antsRegistration’, using brain extracted versions of both T1w reference and the T1w template. The ICBM 152 Nonlinear Asymmetrical template version 2009c was selected for spatial normalization.

### Functional Data Pre-processing

Resting state functional fMRI data were interpolated to 2 x 2 x 2 mm^3^ and underwent spatial smoothing (4 mm FWHM). After smoothing, a standard denoising pipeline was applied [74, 75], which included the regression of potential confounding effects characterized by white matter timeseries (5 CompCor noise components), cerebrospinal fluid timeseries (5 CompCor noise components), motion parameters (XYZ translation and rotation; 6 factors), TRs with excessive motion (129 factors), session and task effects and their first order derivatives (2 factors), QC_cosine regressors (7 components), and linear trends (2 factors) within each functional run. The residual time series data were bandpass frequency filtered between 0.008 Hz and 0.09 Hz [76]. CompCor [77, 78] noise components within white matter and cerebrospinal fluid were estimated by computing the average BOLD signal as well as the largest principal components orthogonal to the BOLD average, motion parameters, and outlier scans within each subject’s eroded segmentation masks. From the number of noise terms included in this denoising strategy, the effective degrees of freedom of the BOLD signal after denoising were estimated to range from 16.2 to 72.9 (average 44.8) across all subjects [79].

For each run of fMRI data, a reference volume and its skull-stripped version were generated using fMRIPrep. Head-motion parameters with respect to the BOLD reference (transformation matrices, and six corresponding rotation and translation parameters) were estimated before spatiotemporal filtering using ‘MCFLIRT’ from FSL. BOLD runs were slice-time corrected using ‘3dTshift’ from AFNI. The BOLD time-series were then resampled onto their original, native space by applying the transforms to correct for head-motion. The BOLD reference was then co-registered to the T1w reference using ‘mri_coreg’ (FreeSurfer) followed by ‘FLIRT’ (FSL) with the boundary-based registration cost-function. Co-registration was configured with twelve degrees of freedom to account for distortions remaining in the BOLD reference. The BOLD time-series were resampled into standard space, generating a pre-processed BOLD run in MNI152nLin2009cAsym space. All resamplings can be performed with a single interpolation step by composing all the pertinent transformations (i.e., head-motion transform matrices, susceptibility distortion correction when available, and co-registrations to anatomical and output spaces). Gridded (volumetric) resamplings were performed using ‘antsApplyTransforms’ (ANTs), configured with Lanczos interpolation to minimize the smoothing effects of other kernels.

Following pre-processing in fMRIPrep, fMRI and T1 anatomical data were analyzed with the BrainVoyager software package (Version 22.4) and in-house scripts drawing on the NeuroElf toolbox written in MATLAB. All functional data underwent temporal high-pass filtering (cutoff: 2 cycles per time course within each run), were smoothed at 6 mm FWHM, and were interpolated to 3 x 3 x 3 mm voxels. A participant-specific general linear model was used to fit beta estimates to the experimental events of interest. Experimental events were convolved with a standard 2-gamma hemodynamic response function. The first derivatives of 3D motion correction from each run were added to all models as regressors of no interest to attract variance attributable to head movement (XYZ translation and rotation; 6 covariates).

### Resting State Functional Connectivity and Region-of-Interest (ROI) Definition

Functional connectivity was computed in the 107 aged-matched neurotypical participants and separately in the 55 non-age-matched neurotypical participants. One participant in the age-matched group was dropped from the analysis due to missing data. Seed-based connectivity maps were estimated characterizing the spatial pattern of functional connectivity with the dPCSA and vPCSA regions. Functional connectivity strength was represented by beta values from a weighted general linear model [80] quantifying the correlation between each seed ROI’s time series to the time series of each voxel in the brain.

MNI coordinates for the dPCSA and vPCSA were extracted using coordinates from Rong and colleagues [81]. In that study, the dPCSA ROI was defined by identifying the peak voxel in left precentral gyrus responding to the ‘Listen + Rehearsal’ condition relative to the ‘Listen + Rest’ condition. The vPCSA ROI was defined by identifying the peak voxel in left precentral gyrus responding to the ‘Listen + Rehearsal’ condition relative to the ‘Resting’ condition. Here, spherical ROIs 5 mm in diameter were drawn around each peak (see Supplementary Figure 1A).

We used the search terms “superior temporal” and “supramarginal” in Neurosynth to independently identify the superior temporal gyrus and anterior supramarginal gyrus, respectively. The superior temporal gyrus (STG) was chosen as an ROI given its association with pitch processing [18, 82–84]; the anterior supramarginal gyrus (aSMG) was chosen as an ROI given its association with somatosensory processing during speech production [27, 40]. Spherical ROIs 5 mm in diameter were then drawn around these masks (see Supplementary Figure 1B).

We computed whole-brain functional connectivity for the seed ROIs (dPCSA, vPCSA), extracted values using the mask ROIs (STG, SMG), and took the mean of all voxels in each mask for each neurotypical participant. A mixed ANOVA with the factors Group (age-matched neurotypical group, non-age-matched neurotypical group), Seed ROIs (dPCSA, vPCSA) and Mask ROIs (STG, aSMG) was then performed.

### Functional MRI Data Extraction and Voxel-based Lesion Activity Mapping

The contrast of ‘Sentence Repetition > Sentence Listen’ was computed in each participant; we then took the mean of all contrast-weighted *t*-values in the dPCSA and vPCSA ROIs, which served as independent variables in voxel-based lesion-activity mapping (VLAM) analyses of sentence repetition. Those same ROIs were used to extract *t*-values from the contrast of ‘Melody Humming > Melody Listen’. The mean of voxels in each ROI served as additional independent variables in VLAM analyses of melody humming.

VLAM uses functional neural responses in an ROI to predict lesion overlap throughout the brain [30]. VLAM was performed using the SVR-LSM toolbox using a univariate model [85]. Only voxels lesioned in at least 10% of participants were included. We controlled for variability in lesion volume using ‘Direct Total Lesion Volume Control’, which divides lesioned voxels in each participant’s lesion map by the square root of the total lesion volume. This correction method places a greater emphasis on smaller lesions when controlling for lesion volume in each participant’s lesion map. Voxelwise statistical significance was determined using a permutation analysis in which the functional neural responses were randomly assigned to a lesion map, and the same procedure as described above was iterated 10,000 times. Voxelwise z-scores were computed for the true data in relation to the mean and standard deviation of voxel-by-voxel null distributions; the resulting z-score map was set to a threshold of z < −1.65 (*p* < .05, one-tailed) to determine chance-level likelihood of identifying a significant effect in each voxel. In all subsequent VLAM analyses, if the resulting voxelwise clusters did not survive permutation analysis we removed clusters with fewer than 500 contiguous voxels before interpreting the results (see [71–73, 86, 87]). The Harvard-Oxford Cortical Probabilistic Atlas assessed the location of significant voxels resulting from each VLAM analysis.

## Data availability

The whole-brain maps of contrast-weighted *t*-values, lesions in MNI space, and analysis code will be made available via a publicly accessible repository upon publication of the manuscript. Participants did not consent to public archival of the MRI data. Researchers interested in accessing the raw MRI data should contact the corresponding author to establish a data use agreement.

## Acknowledgments

This research was supported by the University of Rochester CTSA awards NIH UL TR002001 and NIH KL2 TR001999 from the National Center for Advancing Translational Sciences, and from the Harry W. Fischer fund within the Department of Imaging Sciences at the University of Rochester Medical Center to F.E.G. This research was also supported by NIH grants R21NS076176 and R01NS089069 and NSF grant BCS-1 349 042 to B.Z.M., by a core grant to the Center for Visual Science (P30 EY001319), and by funding support to the Department of Neurosurgery at the University of Rochester by Norman and Arlene Leenhouts. W.B. was supported by the SKAWA Foundation UR Medical Student Fellowship Grant.

## Footnotes

1 Alario and colleagues (2006) note that in humans, the SMA can be anatomically separated from the pre-SMA in the coronal plane at the anterior commissure, which roughly maps to a Y-coordinate of 0 (i.e., when viewing the anterior commissure in the coronal plane, the SMA is posterior to Y = 0, and the pre-SMA is anterior to Y = 0). Although the Harvard-Oxford atlas does not have separate SMA and pre-SMA parcels, the peak Y-coordinate found in our VLAM analysis of sentence repetition in the vPCSA was 2, suggesting this area is the pre-SMA.

