## Supplemental online materials for "Causal Lesion Evidence for Two Motor Speech Coordination Networks in the Brain"

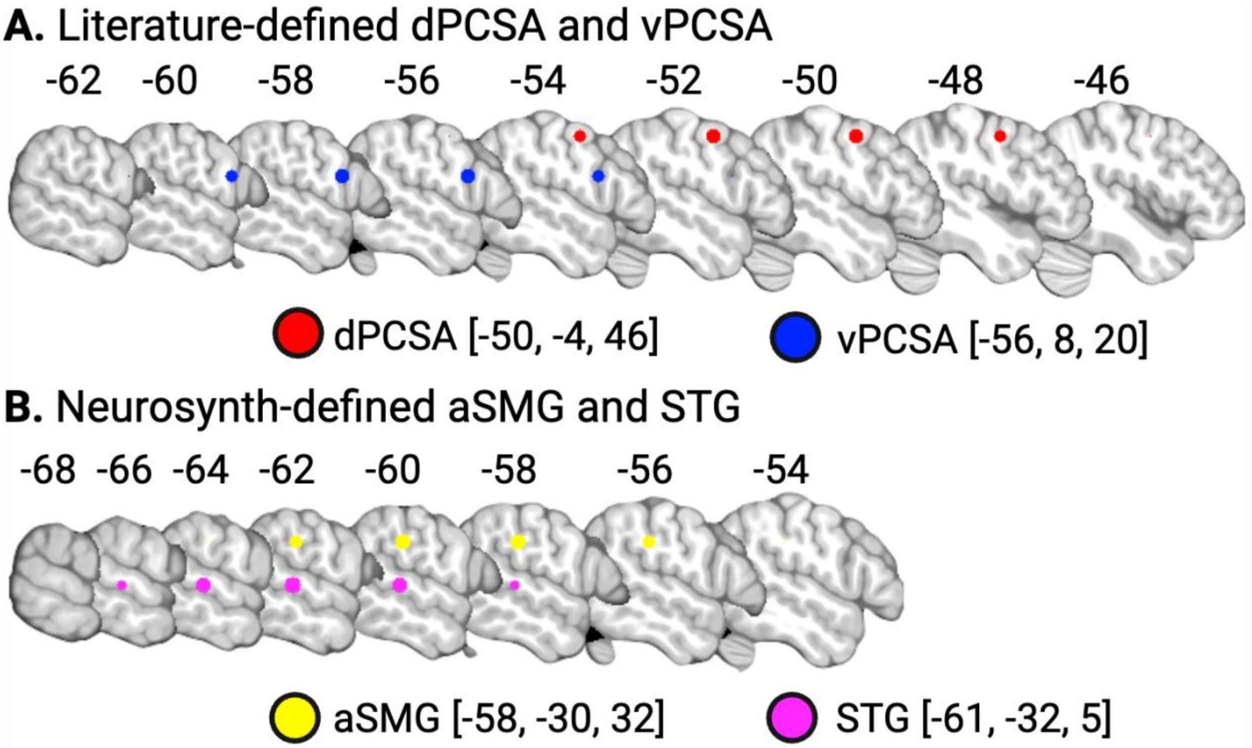


**Supplemental Figure 1. Regions-of-interest.** X-coordinates in MNI space are provided above each sagittal image. **(A)** Spherical ROI representations of the dPCSA and vPCSA derived from Rong and colleagues (2018). These ROIs were used (1) as seed regions in the resting state functional connectivity analysis (see Figure 1) and (2) in the VLAM analyses by extracting contrast-weighted *t*-values separately for sentence and melody listening phases as well as for sentence and melody reproduction phases (see Figure 2). **(B)** MNI coordinates for the superior temporal gyrus (STG) and anterior supramarginal gyrus (aSMG) derived from Neurosynth using the search terms "superior temporal" and "supramarginal," respectively. The automated meta-analysis for the term "superior temporal" included 1422 studies and 50978 activations, and the automated meta-analysis for the term "supramarginal" included 375 studies and 13835 activations. These ROIs were used as mask regions in the resting state functional connectivity analysis (see Figure 1).

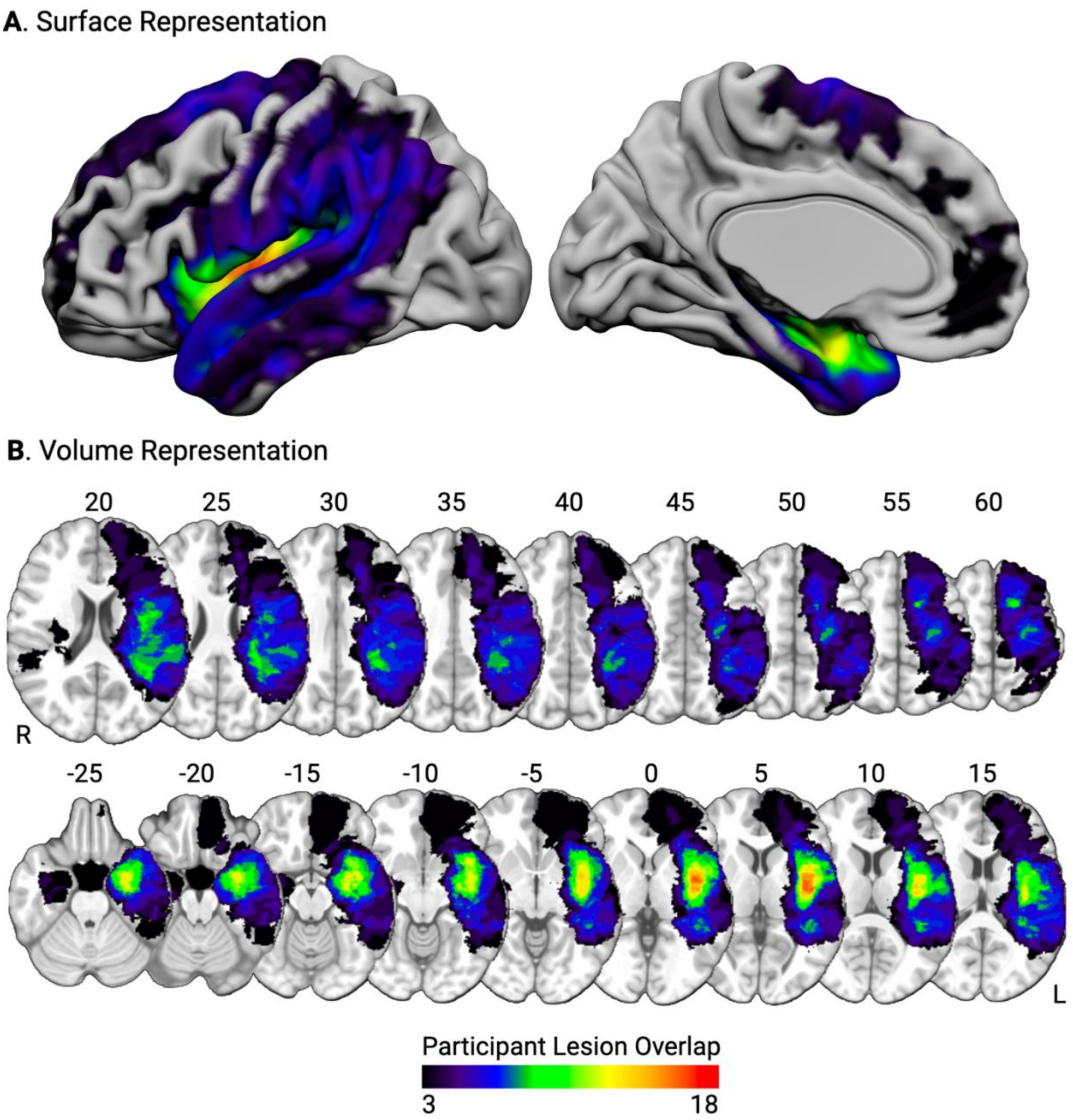


**Supplemental Figure 2. Voxelwise lesion overlap among participants.** Surface-based (A) and volumetric (B; MNI Z coordinate listed above axial slices) renderings of the lesion overlap in voxels among participants. Warmer colors indicate greater lesion overlap among participants.

**Supplementary Table 1.** Demographic information for each participant. *Abbreviations*. Arteriovenous malformation, AVM; Cerebral cavernous malformation, CCM; Dysembryoplastic neuroepithelial tumor, DNET; Glioblastoma multiforme, GBM; Metastatic Carcinoma, Met. Carcinoma; dPCSA, dorsal precentral speech area; vPCSA, ventral precentral speech area; SR > SL, fMRI contrast of 'Sentence Repetition > Sentence Listening'; MH > ML, fMRI contrast of 'Melody Humming > Melody Listening'.

| Participant ID | Age | Diagnosis | Gender | Lesion Volume | Hemisphere | dPCSA SR > SL | dPCSA MH > ML | vPCSA SR > SL | vPCSA MH > ML |
|---|---|---|---|---|---|---|---|---|---|
| 1 | 26 | Astrocytoma | Male | 14309 | Right | 0.56 | 1.3 | -0.14 | 1.23 |
| 2 | 65 | Astrocytoma | Male | 34734 | Left | 7.35 | 5.35 | 1.25 | 0.49 |
| 3 | 71 | Astrocytoma | Male | 19704 | Left | 0.06 | 0.83 | 2.22 | 2.62 |
| 4 | 36 | Astrocytoma | Female | 64557 | Left | 5.09 | 3.9 | 5.76 | 3.24 |
| 5 | 24 | Glioma | Female | 9230 | Left | 0.01 | 0.13 | -0.63 | -2.22 |
| 6 | 25 | CCM | Female | 3993 | Left | -0.02 | 0.31 | 1.45 | 1.7 |
| 7 | 26 | Gliosis | Male | 17726 | Left | -2.24 | -0.53 | -3.8 | 0.02 |
| 8 | 37 | Gliosis | Female | 183 | Left | 0.22 | 0.73 | -1.18 | -1.36 |
| 9 | 28 | GBM | Male | 193209 | Left | -0.22 | -0.3 | 0.19 | -0.51 |
| 10 | 65 | Astrocytoma | Male | 29215 | Right | 0.85 | -0.49 | 2.85 | -0.95 |
| 11 | 61 | GBM | Female | 133252 | Right | -0.92 | -1.91 | 0.25 | -1.3 |
| 12 | 36 | Oligodendroglioma | Female | 56682 | Right | 0.14 | -3.99 | -0.83 | -3.99 |
| 13 | 36 | GBM | Male | 90810 | Right | 4.31 | 4.53 | -1.16 | 0.35 |
| 14 | 65 | GBM | Female | 102939 | Right | -0.4 | 7.16 | 0.47 | -1.17 |
| 15 | 79 | GBM | Female | 35746 | Left | -3.27 | 0.35 | -2.02 | 0.08 |
| 16 | 58 | GBM | Female | 188060 | Left | -3.49 | -3.05 | -0.44 | -1.27 |
| 17 | 45 | Astrocytoma | Male | 114420 | Left | 2.94 | 1.39 | 0.95 | -0.67 |
| 18 | 55 | Astrocytoma | Female | 15297 | Left | -1.45 | -3.12 | 0.2 | -5.8 |
| 19 | 47 | Oligodendroglioma | Male | 99649 | Left | 3.42 | -4.1 | 4.11 | -0.06 |
| 20 | 31 | Oligodendroglioma | Male | 69264 | Left | -2.17 | -1.52 | -2.39 | -2.14 |
| 21 | 51 | Astrocytoma | Female | 36433 | Right | -0.66 | 1.14 | -2.59 | -0.74 |
| 22 | 75 | GBM | Female | 60841 | Left | -0.42 | -3.27 | 1.07 | -1.11 |
| 23 | 42 | GBM | Male | 218617 | Left | -4.86 | -5.92 | -5.94 | -5.36 |
| 24 | 45 | GBM | Female | 276913 | Left | 0.01 | 1.61 | 0.33 | 2.49 |
| 25 | 76 | GBM | Male | 1405 | Left | -0.78 | -1.68 | -4.19 | -6.38 |

| | | | | | | | | | |
|---|---|---|---|---|---|---|---|---|---|
| 26 | 34 | Glioma | Male | 83636 | Left | 3.41 | 0.48 | 0.39 | 1.76 |
| 27 | 67 | Oligodendroglioma | Male | 42216 | Left | 0.93 | 0.45 | -1.19 | -0.53 |
| 28 | 40 | AVM | Male | 31769 | Right | 2.55 | 1.98 | 2.8 | 2.25 |
| 29 | 59 | Astrocytoma | Female | 11939 | Right | 2.32 | 1.92 | 8.27 | -1.06 |
| 30 | 28 | Oligodendroglioma | Male | 43634 | Left | 2.26 | 0.13 | -1.79 | -1.88 |
| 31 | 38 | AVM | Male | 19544 | Left | 1.47 | 2.16 | 1 | -0.96 |
| 32 | 62 | Oligodendroglioma | Female | 25137 | Left | 1.87 | 0.11 | 4.62 | 1.18 |
| 33 | 55 | Astrocytoma | Female | 77928 | Right | 0.79 | 0.9 | 0.72 | 0.62 |
| 34 | 46 | DNET | Female | 10291 | Left | 0.61 | 0.23 | 0.63 | 1.26 |
| 35 | 57 | GBM | Male | 145300 | Left | 1.15 | 1.12 | 1.82 | 1.23 |
| 36 | 67 | GBM | Male | 18604 | Left | 2.34 | -1.42 | 2.13 | -0.39 |
| 37 | 18 | Astrocytoma | Male | 5641 | Right | 1.6 | -0.95 | -0.69 | -2.11 |
| 38 | 46 | Oligodendroglioma | Male | 67034 | Right | -0.53 | -1.54 | -0.91 | -1.53 |
| 39 | 18 | Glioma | Female | 21027 | Left | 0.56 | 1.04 | 0.5 | 0.84 |
| 40 | 58 | Met. Carcinoma | Female | 85172 | Left | -1.5 | 1.71 | -1.25 | 0.01 |
| 41 | 70 | GBM | Female | 38524 | Right | -0.61 | -0.93 | -0.79 | 0.76 |
| 42 | 75 | Astrocytoma | Female | 25094 | Right | -0.38 | 0.07 | -0.53 | -1.27 |
| 43 | 46 | DNET | Female | 34142 | Left | 0.29 | -0.46 | 1.49 | -0.49 |
| 44 | 29 | Astrocytoma | Female | 69361 | Left | 0.72 | -1.39 | -0.51 | -1.27 |
| 45 | 65 | AVM | Male | 35044 | Left | 1.46 | 0.41 | 2.14 | 1.69 |
| 46 | 73 | Glioma | Male | 11361 | Left | -1.89 | -3.13 | 0.25 | -1.63 |
| 47 | 63 | GBM | Male | 16875 | Left | -1.54 | -2.57 | 3.59 | 3.55 |
| 48 | 59 | GBM | Female | 59945 | Left | 0.71 | 0.76 | -0.68 | 0.28 |
| 49 | 35 | Astrocytoma | Male | 47252 | Left | 1.03 | 0.99 | -0.77 | -1.94 |
| 50 | 40 | Astrocytoma | Female | 26814 | Left | 0.5 | -1.12 | 1.63 | -1.72 |
| 51 | 61 | GBM | Male | 83984 | Left | -0.39 | -0.78 | -0.51 | -1.84 |
| 52 | 27 | CCM | Male | 9046 | Right | -0.99 | -1.47 | 0.9 | -0.99 |
| 53 | 44 | Glioma | Male | 14179 | Left | -0.37 | -1.19 | -0.92 | -1.74 |
| 54 | 49 | Glioma | Female | 23800 | Left | 0.13 | 0.86 | 0.64 | -0.57 |
| 55 | 23 | Glioma | Male | 3871 | Right | 1.33 | -0.2 | 1.33 | 1.59 |
| 56 | 23 | Astrocytoma | Male | 30201 | Left | -0.99 | 1.29 | -3 | 1.19 |
| 57 | 32 | Biopsy not performed | Female | 7825 | Left | 0.29 | 1.83 | -2.22 | 0.51 |

| | | | | | | | | | |
|---|---|---|---|---|---|---|---|---|---|
| 58 | 61 | Biopsy not performed | Male | 666 | Right | -1.06 | -0.68 | -3.57 | -2.84 |
| 59 | 30 | Oligodendroglioma | Male | 220690 | Left | 2.5 | 0.82 | 0.77 | 0.39 |
| 60 | 41 | GBM | Female | 51594 | Left | 1.29 | 3.08 | 0.17 | -0.35 |
| 61 | 36 | Astrocytoma | Female | 14672 | Right | -0.54 | 0.94 | 0.38 | 0.34 |
| 62 | 34 | Neurocytoma | Female | 16970 | Left | 0.03 | 0.39 | 0.2 | 1.63 |
| 63 | 22 | Astrocytoma | Female | 23965 | Left | -0.34 | -2.98 | -1.92 | -2.11 |
| 64 | 59 | Meningioma | Female | 80075 | Left | 2.07 | 2.05 | 0.4 | 1.79 |
| 65 | 33 | Biopsy not performed | Female | 16135 | Left | -0.43 | 1.58 | -2.42 | 0.4 |
| 66 | 57 | GBM | Male | 68487 | Left | 0.21 | 2.65 | -1.18 | 1.55 |

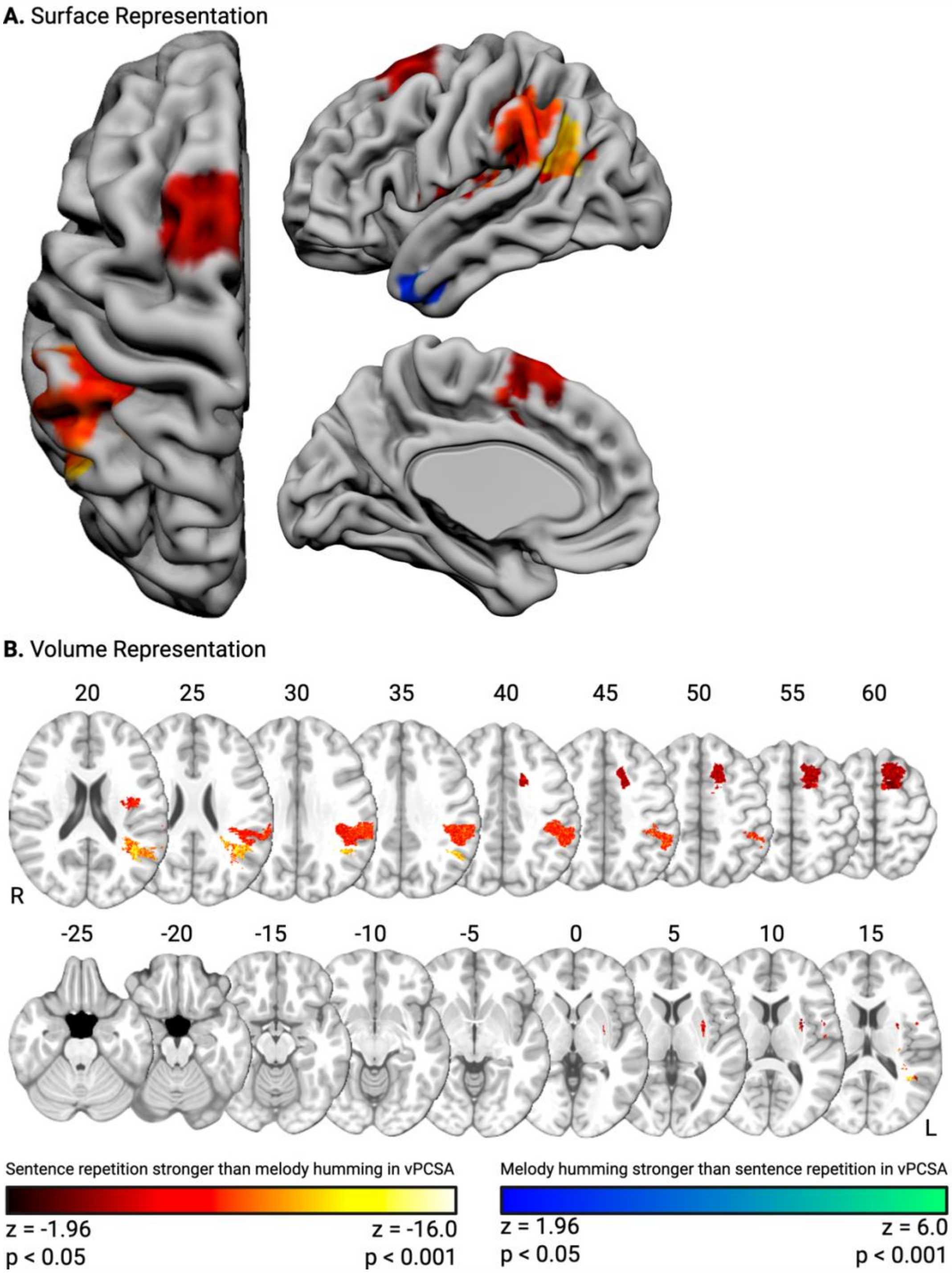


**Supplementary Figure 3. Comparison of VLAM results for vPCSA**. We inspected where the VLAM result of sentence repetition was statistically stronger than the VLAM result of melody humming. Voxelwise values in each map are z-scores; thus, we performed a subtraction analysis in each voxel to quantify the difference between the VLAM result for sentence repetition in the vPCSA subtracting out the VLAM result for melody humming in the vPCSA. A 1,000 iteration permutation analysis then followed, in which the VLAM maps were scrambled prior to subtracting the VLAM result for melody humming from the VLAM result for sentence repetition. The voxelwise difference scores of the true data were then z-scored against the mean permutation-derived voxelwise difference scores. Surface-based (A) and volumetric (B; MNI Z coordinate listed above axial slices) renderings of the lesion sites demonstrated a differentially stronger VLAM effect for sentence repetition (red-to-yellow) and melody humming (blue-to-green). This analysis confirms that the VLAM result in supramarginal gyrus for sentence repetition was statistically stronger than the VLAM result in the supramarginal gyrus for melody humming.

**Supplementary Table 2.** (A) Regions which demonstrated a differentially stronger VLAM effect for sentence repetition in the vPCSA and (B) regions which demonstrated a differentially stronger VLAM effect for melody humming in the vPCSA.

| Region Name | Region Number | Peak MNI Coordinate | | | Peak Z-Score | Percent of Region with Significant Voxels |
|---|---|---|---|---|---|---|
| | | X | Y | Z | | |
| **A. VLAM of sentence repetition in stronger than VLAM of melody humming** | | | | | | |
| Angular gyrus | 41 | -42 | -58 | 25 | -19.68 | 15.81 |
| Anterior supramarginal gyrus | 37 | -62 | -41 | 32 | -16.74 | 40.38 |
| Central operculum | 83 | -37 | -7 | 18 | -15.37 | 4.47 |
| Heschl's gyrus | 89 | -44 | -21 | 11 | -14.85 | 0.60 |
| Posterior supramarginal gyrus | 39 | -48 | -42 | 50 | -14.84 | 14.86 |
| Postcentral gyrus | 33 | -37 | -33 | 42 | -14.51 | 1.33 |
| Superior parietal lobe | 35 | -45 | -41 | 50 | -13.00 | 0.77 |
| Parietal operculum | 85 | -48 | -41 | 24 | -12.36 | 17.58 |
| Superior frontal gyrus | 5 | -20 | 4 | 53 | -10.75 | 12.85 |
| Insular cortex | 3 | -35 | -10 | 18 | -10.61 | 0.28 |
| Supplementary motor area | 51 | -11 | -3 | 53 | -10.00 | 3.66 |
| Paracingulate gyrus | 55 | -11 | 15 | 46 | -9.84 | 0.48 |
| Superior lateral occipital cortex | 43 | -43 | -68 | 25 | -8.15 | <0.01 |
| Precentral gyrus | 13 | -51 | 4 | 9 | -5.95 | 0.02 |
| **B. VLAM of melody humming stronger than VLAM of sentence repetition** | | | | | | |
| Temporal Pole | 15 | -50 | 8 | -31 | 4.99 | 1.50 |
| Anterior middle temporal gyrus | 21 | -48 | 0 | -31 | 2.40 | 0.03 |
| Anterior inferior temporal gyrus | 27 | -46 | 1 | -35 | 2.94 | 0.20 |
